# Altered EEG markers of reward learning during abstinence in alcohol dependence: a probabilistic reversal learning study

**DOI:** 10.1101/2024.11.03.620234

**Authors:** Mica Komarnyckyj, Chris Retzler, Anna Murphy, Ioannis Delis, Elsa Fouragnan

**Affiliations:** Division of Psychology & Mental Health, University of Manchester, Oxford Rd, Manchester, M13 9PL; Centre for Cognition and Neuroscience, University of Huddersfield, Queensgate, Huddersfield, HD1 3DH, UK; School of Biomedical Sciences, University of Leeds, West Yorkshire, LS2 9JT, UK; School of Psychology, University of Plymouth, Plymouth, Portland Square, PL4 8AA, UK; Brain Research Imaging Centre, Faculty of Health, University of Plymouth, Plymouth PL6 8BU, UK

## Abstract

Maladaptive reward learning and decision-making circuity are key factors in the onset and progression of alcohol use disorder and have therefore emerged as key targets for neuropsychological and pharmacological interventions. Probabilistic reversal learning studies have consistently reported impaired learning in recently detoxified alcohol dependent (AD) participants. However, the neural and behavioural changes associated with reward learning which occur throughout abstinence remain unexplored. Here, we show that AD participants, with mean abstinence of 20 months, exhibit intact behavioural performance within an electroencephalography (EEG) probabilistic reversal learning task. Reinforcement learning modelling reveals reward and punishment related learning rates and exploration rates are comparable between AD and healthy control (HC) participants, suggesting recovery of even the nuanced aspects of learning in longer term abstinence. However, EEG analysis indicates that AD, compared to HC participants, show globally elevated event-related potential (ERP) feedback related negativity (FRN) following reward valuation. Furthermore, Feedback-P3 valence prediction error signal is negatively associated with abstinence duration indicating a potential state marker of AD recovery. We then employ unsupervised machine learning (canonical polyadic tensor decomposition) to identify spatiotemporal EEG patterns of reward valuation in a purely data-driven manner. Classification analysis shows these tensor components can predict group membership with 80.4% accuracy. By probing group differences in tensor components, we discover early hyperfunctioning in centro-frontal regions linked to alcohol dependence and associated with early abstinence. The clinically meaningful EEG biomarkers presented here could guide the development of more targeted treatments and support big data approaches to objective patient monitoring.

## INTRODUCTION

Alcohol dependency is characterised by repeating cycles of craving, uncontrollable consumption, increased tolerance, withdrawal and relapse^1,2^. A core symptom of alcohol dependency is an inability to stop drinking despite harmful consequences to health, safety, relationships, and capacity to work^3^. This failure to learn from bad experiences and change future behaviour to avoid the negative impacts of drinking, are thought to be mediated by dysfunctional learning and decision-making brain circuits^4–6^.

The probabilistic reversal learning task (PRLT) disentangles important components of decision-making and learning which are maladaptive in alcohol dependency, including encoding, valuation and evaluating decision outcome through prediction error (difference between expected and received rewards)^6–10^ (Fig. 1A). This error signal tracks how individuals adjust their strategies to changing reward probabilities and can be extracted through reinforcement learning models. Importantly, the PRLT offers insights into the activity of neural nodes and circuits involved in reward learning, feedback processing and cognitive flexibility^11,12^. Functional magnetic resonance imaging (fMRI) studies using the PRLT have demonstrated correlations between prediction error and brain areas involved in reward processing, including the striatum and prefrontal cortex subregions in healthy individuals. This relationship has been shown to be dysfunctional in alcohol dependent (AD) people, particularly through abnormal connectivity between the dorsolateral prefrontal cortex (DLPFC) and striatum which predicts impaired learning from rewards^6,9^.

**Fig. 1.**
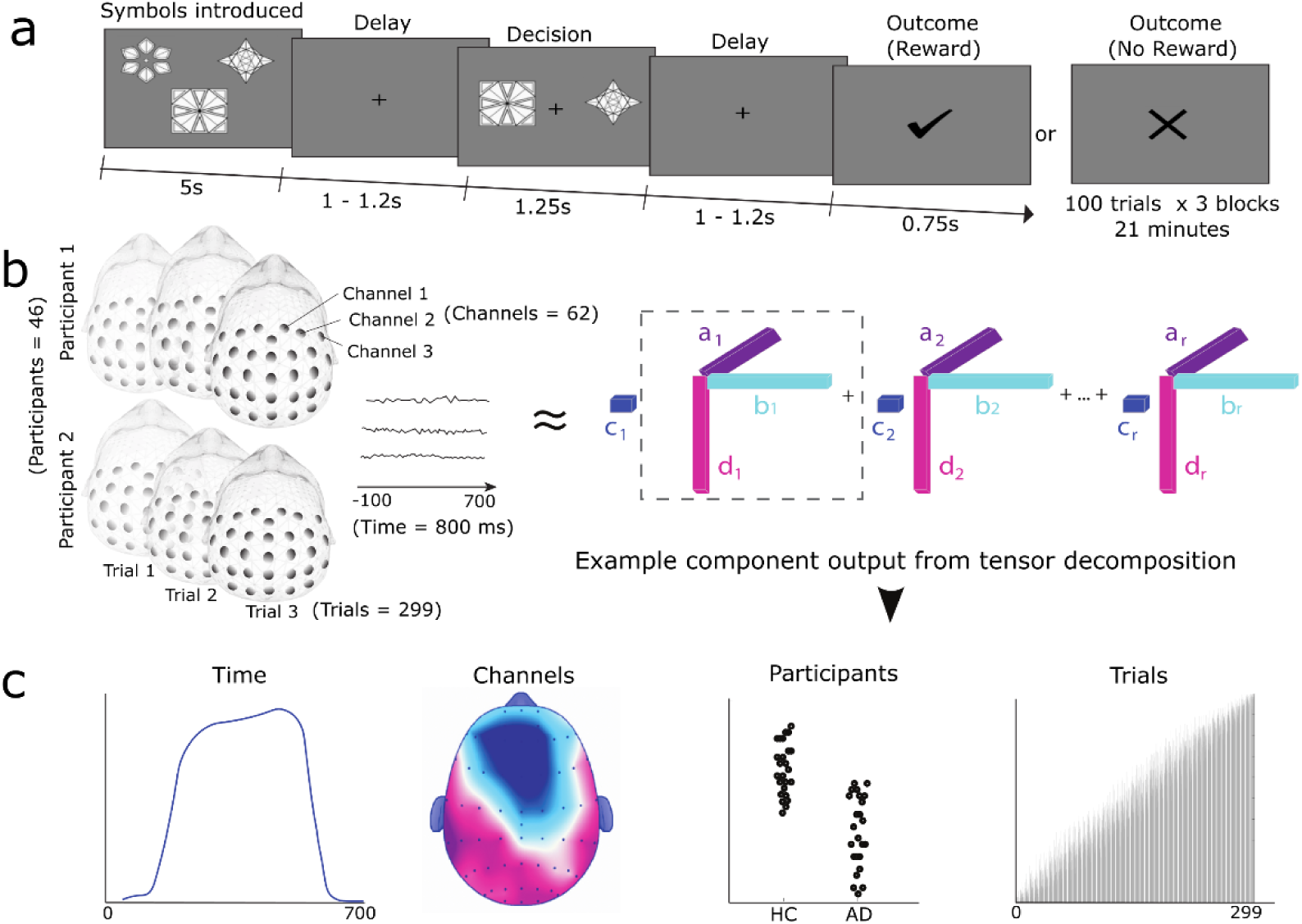
Overview of the reversal learning task and tensor decomposition. **(a)** Diagram of reversal learning task. At the start of each block a set of three abstract symbols was shown to familiarise participants with the symbols before the trials began. Every trial began with a jittered delay (1 – 1.2 s) followed by two abstract symbols (selected from the set of three) which were shown for 1.25 s. During this time, participants pressed the left or the right arrow on the keyboard to select which of the two symbols (left or right) they believed was the high reward probability symbol and therefore more likely to lead to a reward outcome (i.e., tick). The fixation cross flickered for 100 ms when a selection was made. Finally, the outcome (tick or cross) was shown for 0.75 s after a second jittered delay. **(b)** Schematic representation of tensor decomposition. The entire 4-dimensional EEG data (Spatial (i.e., 62 channels) × Temporal (i.e., 401 points = 500 ms) × Participants (n = 46) × Trials (n = 299) (left) is fed into the non-negative canonical polyadic tensor decomposition algorithm <u>X</u> ∼ A ◦ B ◦ C ◦ D ◦ E (right) resulting in 4-dimensional tensor components. **(c)** Example component from tensor decomposition. From left to right, each resulting tensor component includes: a temporal factor, reflecting time points across the trials during which component recruitment peaked; a spatial factor, reflecting the strength of component activation across the 62-channels (i.e., EEG scalp topography); a participant factor, illustrating how strongly individual participants recruited the component; and a trials factor, showing how recruitment of the component changed across the course of the task.

Despite electroencephalography (EEG) being a powerful tool for capturing neural activity during decision-making with high temporal resolution^11–13^, no studies have combined it with the PRLT to explore alcohol use disorder (AUD) (including at-risk, problem drinkers or dependent populations). Here, we present the first study of this kind, investigating EEG reward learning and decision-making signals in an abstinent AD population. Compared with fMRI, EEG offers enhanced temporal resolution, lower cost, improved portability and patient tolerance: significant advantages in translational applications targeting maladaptive decision-making circuitry in addiction^14,15^. Prior research has shown the value of EEG in identifying those at-risk of AUD^16–18^, predicting AUD maintenance and remission^19^, and evaluating novel treatments^20–23^.

Here, we applied a computational reinforcement learning model to explore hidden behavioural components of the task, such as learning rate and exploration^7^. We hypothesised impaired learning performance in AD compared to HC participants, since prior PRLT studies have shown alcohol dependency is associated with making fewer correct choices^6,8^, achieving fewer learning reversals^6,10^ and lower learning rates^9^. Event-related potential (ERP) analyses focussed on the feedback related negativity (FRN) and feedback-P3, identified as key timepoints in adaptive decision-making^13^ and potential biomarkers of AUD^17,21,24,25^. Based on previous studies of reward valuation, we hypothesised decreased FRN amplitudes in AD compared to HC participants^17,25,26^, however our investigation of the feedback-P3 was exploratory given the previous findings are mixed^17,25^.

ERP is the one of the most common methods of EEG analysis, however by focusing on pre-determined time windows and electrodes, a large portion of data is often overlooked and there is an increased risk of selective reporting which can inflate Type I error^27^. Additionally, ERP waveforms may reflect multiple neural processes which overlap in time and are challenging to disentangle. To overcome these limitations, we applied a data-driven tensor decomposition algorithm (non-negative canonical polyadic decomposition)^28^ to the entire EEG dataset (all participants, electrodes, time points, and trials) (Fig. 1B). This unsupervised machine learning method transformed the multidimensional EEG data into simpler tensor components, disentangling overlapping processes that contribute to observed signals, allowing us to identify underlying spatiotemporal components associated with alcohol dependence.

Another novel aspect of this study lies in the diversity of abstinence duration in our AD sample (1–76 months). This allowed investigation of neural and behavioural changes related to abstinence, which has not been possible in prior PRLT research^6,8,9^. Here, we identify dynamic EEG state markers (feedback-P3 and tensor components) that evolve as abstinence duration increases. Our mixed effects models found no associations between the FRN and abstinence or previous drinking behaviours. We propose the decreased FRN amplitudes uncovered here, which are consistently documented in other reward studies over the life course of AUD^17,25,26^, may therefore be a trait marker with potential for predicting those at risk of the disease.

## RESULTS

### Behavioural findings

Within this study 20 abstinent AD participants (mean abstinence of 20 months, range: 1 – 76 months) and 26 HC participants (Table 1) performed a probabilistic reversal learning task whilst 64-channel EEG data were recorded (Fig. 1). Although learning within the PRLT was hypothesised to be abnormal in AD based on a number of prior studies with recently detoxified AD participants^6,9,10^, here we did not find behavioural evidence to support this. There were no group differences in rection time, number of correct choices made (i.e., trials where the high probability symbol was selected), learning reversals (i.e., reflects how many times participants adequately learnt the rule), win-stay trials (i.e., positive outcome trials which were followed by trials in which the same symbol was selected) or lose-switch trials (i.e., negative outcome trials which were followed by trials in which a different symbol was selected) (Table 2). Demonstrating the cognitive demands of the PRLT we found a moderate positive correlation between IQ and learning reversals achieved (Spearman R(44) = .37, *p* = .012).

**Table 1.** Participant demographic and clinical characteristics.

|  | Healthy Control<br>N = 26 | Alcohol<br>Dependent<br>N = 20 | *p* |
| --- | --- | --- | --- |
| **Demographic variables (mean ± SD)** |  |  |  |
| Sex (Male/Female) | 16/10 | 14/6 | 0.989 |
| Age | 40.92 ± 10.50 | 44.70 ± 9.07 | 0.207 |
| IQ (predicted WAIS based on WTAR) | 111.23 ± 4.85 | 108.90 ± 6.37 | 0.166 |
| Smoking status (Smoking/Non-Smoking) | 10/16 | 10/10 | 0.629 |
| Handedness (EHI > 0 / EHI ≤ 0) | | | 0.423 |
| **Clinical variables (mean ± SD)** |  |  |  |
| Age of first alcoholic drink | 16.13 ± 3.40 | 12.50 ± 2.59 | 0.0004 |
| Alcohol units per adult year of drinking | 761 ± 753 | 7714 ± 3007 | < 0.0001 |
| AUDIT | 3.92 ± 4.15 | 34.55 ± 3.44 | < 0.0001 |
| OCDS | 1.35 ± 2.71 | 11.35 ± 9.13 | 0.0001 |
| BDI | 6.04 ± 5.76 | 15.95 ± 12.52 | 0.003 |
| GAD-7 | 2.46 ± 3.31 | 6.00 ± 6.31 | 0.0167 |
| CTQ | 44.31 ± 16.24 | 49.25 ± 28.04 | 0.488 |

Group differences were evaluated using independent t-tests for continuous variables and Chi-squared test for categorical variables (i.e., sex and smoking status). SD = standard deviation; IQ = Intelligence Quotient; WAIS = Wechsler Adult Intelligence Scale; WTAR = Wechsler Test of Adult Reading; IQ = Intelligence Quotient; EHI = Edinburgh handedness inventory score; AUDIT = Alcohol Use Disorder Identification Test; OCDS = Obsessive Compulsive Drinking Scale; Beck's Depression Inventory = BDI; GAD-7 = Generalised Anxiety Disorder Assessment; CTQ = Childhood Trauma Questionnaire).

**Table 2.**
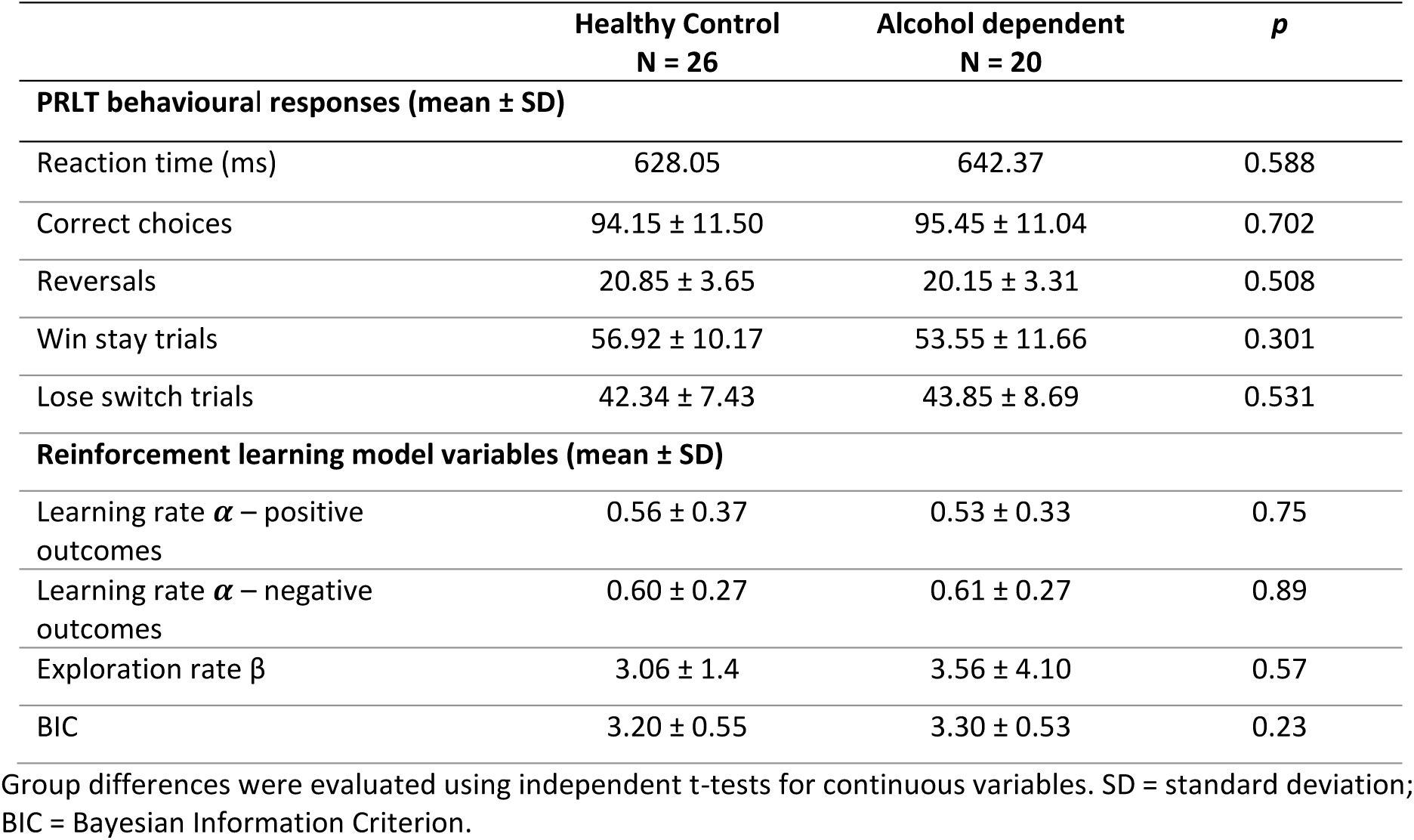
Differences in behavioural and reinforcement learning between HC and AD.

We evaluated three different model-free RL algorithms to estimate trial-wise scaling of PE during the PRLT including a classical Rescorla–Wagner (RW) model, reward-punishment (RP) model (sperate learning rates for positive and negative outcomes) and dynamic learning rate (DRL) model. The RP model results are presented in Table 2 since this model had the best model fit indicated by the lowest BIC (BIC_RW_ = 164.04; BIC_RP_ = 150.93; BIC_DLR_ = 151.99). All variables output from the RP model were found to be equivalent between the HC and AD group, with no differences in the learning rate (*α*) from positive or negative outcomes, exploration rate (β) and model fit (BIC). Further analysis confirmed there were no group differences in these variables when output from the RW and the DLR model.

### FRN amplitude is reduced in alcohol dependence

To investigate valence processing, we conducted a 2×2 ANCOVA (trial outcome: positive-PE, negative-PE; group: HC vs AD) on FRN mean amplitude (180 - 260 ms). For salience processing, we performed a separate 2×2 ANCOVA (trial PE: high vs low-PE; group: HC vs AD) on FRN mean amplitude.

Our ERP findings support the hypothesis of reduced FRN amplitudes in alcohol dependency at time of reward valuation^17,25,26^, we found a significant main effect of group in both the FRN valence ANCOVA (*F*(1, 42) = 5.87, *p* = .020, η_p_^2^ = .123)(Fig. 2A) and magnitude ANCOVA (*F*(1, 42) = 8.17, *p* = .007, η_p_^2^ = .163) (Fig. 2B), with FRN amplitudes significantly smaller (i.e., less negative) for the AD compared to the HC group. The topographical plots (Fig. 2C & 2D) illustrate a broad elevated signal (including central, frontal and parietal regions) at the peak of the FRN (200 – 235 ms), which occurred across all conditions (positive-PE, negative-PE, high-PE and low-PE) in the AD compared to HC group.

**Fig. 2.**
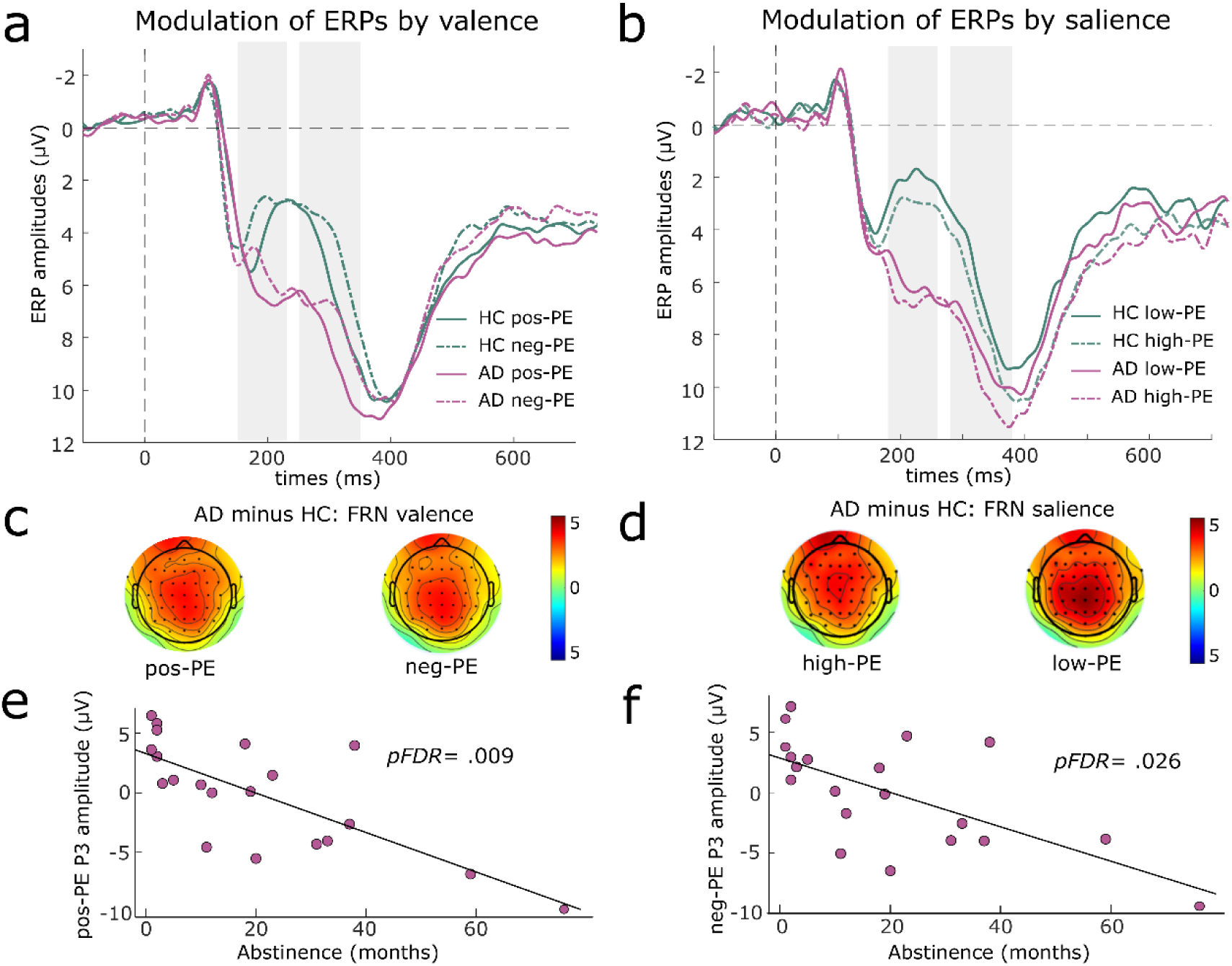
Differences in modulation of ERPs by valence and salience between AD and HC. Average ERP components computed over central electrodes (FC1, FCz, FC2, C1, Cz, C2, CP1, CPz and CP2) for the healthy control (HC) (n = 26) and alcohol dependent (AD) (n = 20) groups. The first and second grey shaded bars depict the time windows for the FRN and feedback-P3 respectively. ERPs are plotted with the negative y-axis pointing up. **(a)** Modulation of ERPs by valence. Negative prediction error (neg-PE) is shown by a dashed line and positive prediction error (pos-PE) by a sold line. **(b)** Modulation of ERPs by salience. Low absolute prediction error (low-PE) is shown by a dashed line and high absolute prediction error (high-PE) by a sold line. **(c)** Scalp topographies are shown for AD minus HC group (illustrating regions of increased activation in AD) at the peak of the FRN (200 – 235 ms), for valence that is pos-PE (left) and neg-PE (right) outcomes. **(d)** Scalp topographies are shown for AD minus HC group at the peak of the FRN (200 – 235 ms), for salience, that is high-PE (left) and low-PE (right) outcomes. **(e)** Partial correlation between feedback-P3 pos-PE amplitude and abstinence duration in months (i.e., linear relationship whilst controlling for the effect of other variables in the mixed effect model including age, IQ, units of alcohol consumed). FDR = False Discovery Rate. **(f)** Partial correlation between feedback-P3 neg-PE amplitude and abstinence duration in months.

There were significantly larger FRN amplitudes for low-PE compared to high-PE outcomes, demonstrated by the significant main effect of prediction error magnitude (*F*(1, 42) = 6.11, *p* = .018, η_p_^2^ = .127). This effect was driven by differences between conditions in the HC group (*t*(42) = 2.12, *p* = .039) which were absent in the AD group (*t*(42) = 1.37, *p* = .179) (Fig. 2B). The main effect of prediction error valence was nonsignificant (*F*(1, 42) = 1.37, *p* = .249, η_p_^2^ = .032) i.e., there was no difference in FRN amplitude between positive-PE vs negative-PE outcomes. The group by condition (valence and magnitude) interaction terms were nonsignificant in the FRN models.

### No group differences in feedback-P3 amplitude

Similarly to the FRN, we conducted a 2×2 ANCOVA (trial outcome: positive-PE, negative-PE; group: HC vs AD) on feedback-P3 mean amplitude (280 - 380 ms). For salience processing, we performed a separate 2×2 ANCOVA (trial PE: high-PE vs low-PE; group: HC vs AD) on feedback-P3 mean amplitude.

The feedback-P3 has been highlighted as a potential ERP biomarker of AUD in prior research ^17,21,24,25^, however the main effect of group did not reach significance in feedback-P3 valence (*F*(1, 42) = 1.32, *p* = .256, η_p_^2^ = .035) and magnitude model (*F*(1, 42) = 2.00, *p* = .164, η_p_^2^ = 0.046). There were significantly larger feedback-P3 amplitudes for positive-PE compared to negative-PE outcomes (Fig. 2A), demonstrated by a significant main effect of prediction error valence (*F*(1, 42) = 26.82, *p* < .001, η_p_^2^ = .390). The feedback-P3 was also modulated by magnitude with significantly larger amplitudes for high-PE compared to low-PE outcomes (Fig. 2B), demonstrated by a significant main effect of prediction error magnitude (*F*(1, 42) = 6.14, *p* = .017, η_p_^2^ = .128). The group by condition (valence and magnitude) interaction terms were nonsignificant in the feedback-P3 models.

### Spatiotemporal dynamics of feedback processing via tensor decomposition

The feedback-locked epochs (i.e., trials) (K) were reduced to the main timepoints (T) of interest for feedback processing (-100 and 700 ms) and then collated across all participants (P) and electrodes (E) into a 4-way data tensor *X*^(4)^ ∈ ℝ ^*T* × *E* × *P* × *K*^, where P = 46, T = 401, E = 62 and K = 299. A tensor decomposition model based on NCP with a hierarchical alternating least squares method was applied to the 4-way data tensor (Fig. 1). The data-driven unsupervised machine learning method uncovered four reward valuation components with overlapping spatial distributions and time windows but distinct temporal peaks (Fig. 3).

**Fig. 3.**
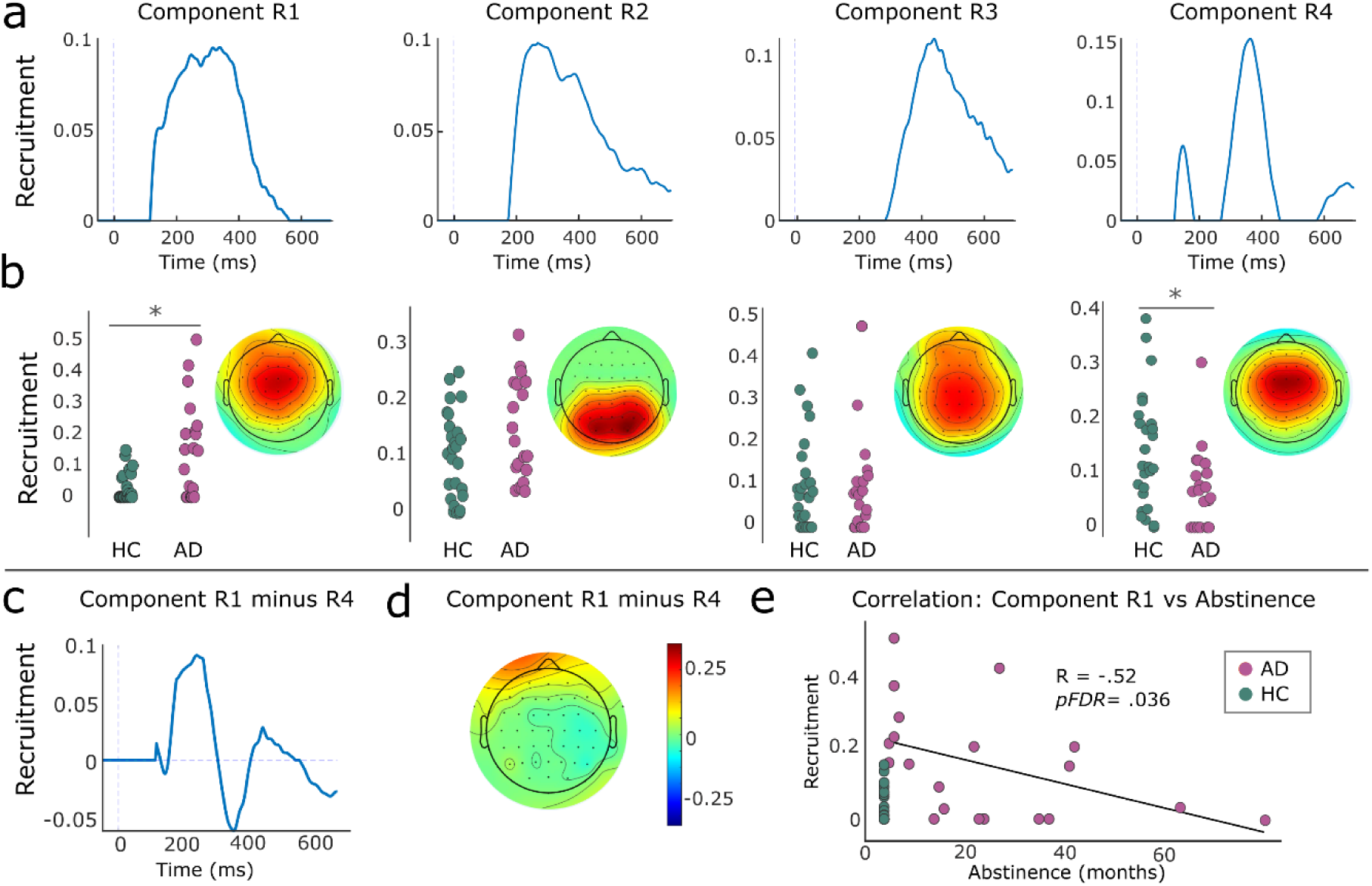
Spatiotemporal recruitment of tensor decomposition composition components. **(a)** Estimated temporal factors from tensor decomposition. The feedback locked epochs were reduced to the main timepoints of interest for feedback processing (-100 and 700 ms). Data were decomposed using NCP into 4 outer product tensor components, Rank (R) = 4. From left to right, temporal factors of the Rank 1 (R1) to Rank 4 (R4) components are shown. **(b)** Estimated participant and spatial factors from tensor decomposition. From left to right, participant factors (including 26 healthy control (HC) and 20 alcohol dependent AD (AD)) (i.e., bar plots) and spatial factors (i.e., EEG scalp topography for 62-channels) of the R1 to R4 components are shown. Significant differences in component recruitment between HC and AD group are shown using a * on the participant factors. **(c)** Temporal factor of component R1 minus R4. **(d)** Spatial factor of component R1 minus R4. **(e)** Spearman correlation between participant factor R1 (most strongly recruited in AD) and abstinence duration in months. FDR = False Discovery Rate.

The first component (R1) occurred across a broad temporal window from 118 ms to 562 ms (Fig. 3A) and had greater recruitment in AD (mean = 0.15, s.d. = 0.15) compared to HC (mean = 0.03, s.d. = 0.05) (*t_44_* = 3.45, *p_FDR_ =* 0.009, 95% CI, CI = [0.69 1.89], Cohen D = 1.15) (Fig. 3B). In contrast, the fourth component (R4) began at 150 ms with a pronounced peak at 366 ms (Fig. 3A) and had greater recruitment in HC (mean = 0.14, s.d. = 0.11) compared to AD (mean = 0.07, s.d. = 0.07) (*t_44_* = -2.45, *p_FDR_ =* 0.023, 95% CI, CI = [-1.38 -0.14], Cohen D = -0.72) (Fig. 3B). By subtracting component R4 from component R1 to identify their spatiotemporal differences (Fig. 3C & Fig. 3D), we observe that they both reflect comparable centro-frontal regions, but component R1 (predominantly activated by AD) begins sooner after feedback (peaking at FRN), whereas component R4 (predominantly activated by HC) starts at a later stage (peaking at feedback-P3).

The second component (r = 2) had a focussed parietal distribution with a temporal peak from 234 ms to 316 ms. The participant factor values for the HC group (mean = 0.10, s.d. = 0.08) and AD group (mean = 0.14, s.d. = 0.09) were not significantly different (*t*(44) = -1.64, *p_FDR_* = 0.155, 95% CI, CI = [-0.08 1.16], Cohen D = 0.49). The third component (r = 3) peaked at 448 ms with a broad centro-fronto-pareital spatial distribution. The participant factor values for the HC group (mean = 0.10, s.d. = 0.11) and AD group (mean = 0.09, s.d. = 0.12) were not significantly different, *t*(44) = 0.18, *p_FDR_* = 0.858, 95% CI, CI = [-0.67 0.55], Cohen D = -0.05).

### Classifying alcohol and control participants from tensor data

We then asked if component activations from tensor decomposition could predict participant group (AD vs HC). We used each participant factor, representing component recruitment at the subject level (Fig. 3B), as a predictor of group (AD positive class, HC negative class) using a medium Gaussian SVM classifier. As expected from the results of the two-sample t-tests, the component R1 resulted in the best classifier performance (Accuracy = 76.1%, AUC = 0.66) when compared to other individual components (Accuracy < 61%) (Supplementary Results). We then tested if combinations of participant factors, extracted from multiple tensor components would increase classification power. We found that using participant factors from R1, R2 and R4 in combination (Fig. 3B) achieved the best classifier performance (Accuracy = 80.4%, AUC = 0.73) (Fig. 4A). This model had a false positive rate of 0%, meaning it did not misclassify any HC participants as AD, and a true positive rate of 55%, meaning 45% of AD were misclassified (Fig. 4B) due to similar component recruitment to HC participants (i.e., illustrated as purple crosses on Fig. 4C and Fig. 4D).

**Fig 4.**
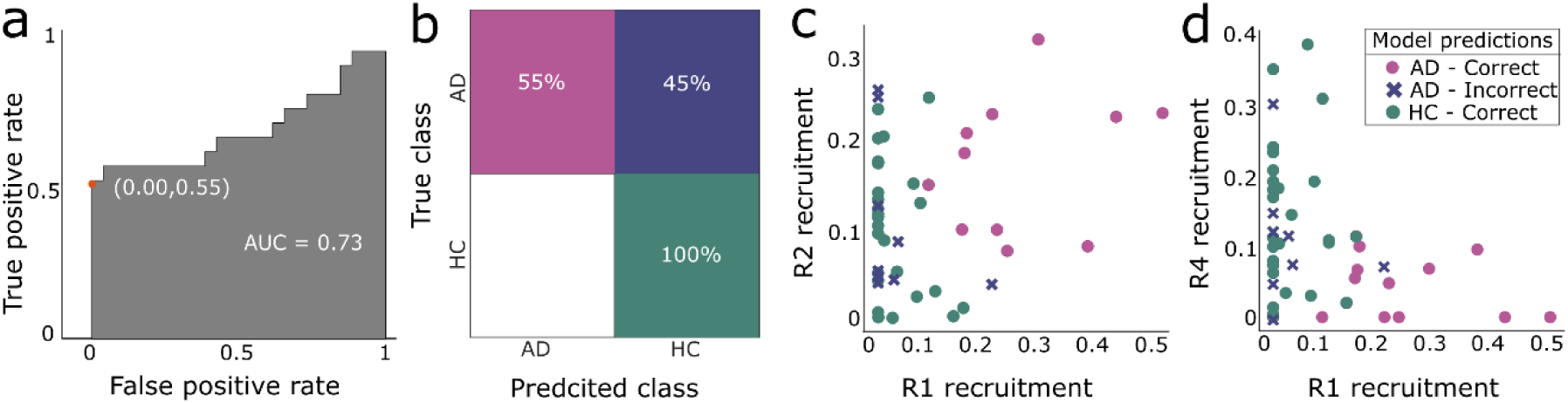
Classification performance for alcohol dependence identification from EEG data. Performance of Medium Gaussian Support Vector Machine (SVM) classifier trained using leave-one-out cross validation, shown for the best performing model which used a combination three participant factors (R1-R2-R4) to predict participant group: alcohol dependent (AD) (n =20) vs healthy control (HC) (n = 26). **(a)** Receiver operating characteristic (ROC) for SVM classifier. **(b)** Confusion matrix for SVM classifier. **(c)** Scatter plot showing classification performance in a two-dimensions based on participants factors from component R1 (x-axis) and component R2 (y-axis). **(d)** Scatter plot showing classification performance in a two-dimensions based participant factors of component R1 (x-axis) and component R4(y-axis).

### Associations with abstinence duration

We used mixed effects linear models to explore whether FRN and feedback-P3 ERP amplitudes could be predicted by clinical severity (including abstinence and alcohol consumption) whilst controlling for age and IQ. We found a main effect of abstinence within the feedback-P3 pos-PE model (*t*_16_ = - 3.98, *p_FDR_ =* 0.009) (Fig. 2E) and feedback-P3 neg-PE model (*t*_16_ = -3.12, *p_FDR_* = 0.026)(Fig. 2F); evidencing that the feedback-P3 amplitude for both positively and negatively valanced feedback is diminished as abstinence duration increases. All other main effects within these models were nonsignificant.

We found a moderate positive correlation between abstinence and learning reversals (Spearman R(18) = .47, *p_FDR_* = .036) and between abstinence and IQ (Spearman R(18) = 0.49, *p_FDR_* = .039), suggesting people who have been abstinent for longer perform better in the task and have improved IQ. Furthermore, we found a moderate negative correlation between recruitment of the first component (R1) and abstinence duration in the AD group (Spearman R(18) = -.52, *p_FDR_* = .036) suggesting longer abstinence is associated with lower recruitment of the component (Fig. 3E).

For the best performing classification model from the tensor decomposition composition analysis (combining R1, R2, R4), we found that correctly classified participants had a shorter abstinence duration (mean = 11.36, range = 1 - 38 months) compared to incorrectly classified participants (mean = 30.89, range = 10 – 76 months) (*t*_18_= -2.35, *p* = .0303, 95% CI, CI = [-2.37 -0.21], Cohen D = -1.06). Notably, all 7 AD participants who had been abstinent for less than 10 months were correctly classified in this model.

## DISCUSSION

We combined hypothesis-driven ERP and data-driven tensor decomposition analyses to uncover EEG state and trait markers of alcohol dependence during a probabilistic reversal learning task. Against our predictions, AD and HC had comparable performance during this learning task but this can be explained by the large variability in abstinence. Using tensor decomposition, we identified spatiotemporal EEG patterns of reward valuation in a purely data-driven manner. By probing group differences in tensor components, we discovered early hyperfunctioning in centro-frontal regions linked to alcohol dependence and associated with early abstinence.

Research has shown that newly abstinent AD populations (under 1.5 months^6,8,9^) have impaired performance on the PRLT. A recent study used a reward-punishment reinforcement learning model similar to the one presented here, demonstrating that AD participants have increased learning from punishment, reduced learning from rewarding feedback and make more random choices^8^. Earlier studies found alcohol dependency was associated with making less correct choices^6^, learning reversals^6,10^ and lower learning rates^9^. Our study did not replicate these findings; we observed equivalent performance and learning between AD and HC. This divergence may be attributed to the fact that our sample had a mean abstinence duration of 20 months. Correlation analyses demonstrated that longer abstinence was associated with improved task performance and IQ of AD participants, consistent with prior research showing impaired executive functions improved in protracted abstinence^29,30^. These findings support the theory that the deficits in reversal learning observed in AD previously^6,8,9^ may be the behavioural state marker which results from chronic alcohol use rather than an underlying trait in alcohol dependent individuals.

The FRN is believed to reflect phasic dopamine activity during reinforcement learning, mirroring reward prediction error signals which arise from dopaminergic projections to the anterior cingulate cortex (ACC)^31–33^. The reduced FRN amplitudes (i.e., more positive amplitudes) shown here in the AD group occur due to globally elevated EEG signal which spans central, frontal and parietal regions (Fig. 2C & 2D) and may reflect dysfunctional reward prediction error signalling originating within the ACC^17,31,34^. Several studies have found similar decreased FRN amplitudes during reward valuation in abstinent AD populations^25,34^ and at-risk drinkers, based on their hazardous drinking^17^, family history^35^ and genetic markers^26^. We found no link between the decreased FRN and prior alcohol use or abstinence (i.e., states of alcohol dependence) suggesting it could be an underlying trait marker for AUD^36^. Importantly, the attenuated FRN appears robust across the life course of AUD (predisposing individuals to the disorder and persisting into their recovery) and is evident using a range of reward paradigms^17,25,26,34,35^. Identifying stable neurophysiological markers such as the FRN, could be invaluable for translational psychiatry, facilitating earlier detection of at-risk individuals and supporting the development of intervention strategies.#

Notably, in contrast to the common reinforcement learning theory that proposes the FRN is enhanced for negative compared to positive feedback^31–33^, here we did not demonstrate FRN sensitivity to reward valence (i.e., positive-PE vs negative-PE). Instead, in the HC group only, the FRN was enhanced for low-PE compared to high-PE outcomes, indicating sensitivity to outcome magnitude, which was absent in the AD group due to the globally elevated signal within the FRN window. Newer theories propose the FRN reflects the magnitude (i.e., salience) of prediction errors, responding to important or unexpected events^32^. Consistent with our findings, previous studies also found enhanced FRN amplitudes for low-magnitude outcomes^37,38^. The FRN’s response to feedback remains debated, possibly due to varying experimental paradigms and methods ^32^.

The feedback-P3 is thought to reflect motivational significance of feedback and attentional allocation during reward valuation. In line with this theory, we found increased feedback-P3 for high-PE (i.e., more salient) compared to low-PE outcomes^32^. Prior research of reward valuation reported detoxified AD have lower feedback-P3 than HC^25^, and conversely at-risk drinkers have elevated feedback-P3 amplitudes compared to low-risk drinkers^17^. Similar to^34^, this study did not find group differences in the feedback-P3 measured at reward valuation for any outcome. We did however find that as abstinence duration increased, feedback-P3 amplitude from pos-PE and neg-PE outcomes, decreased. This provides some evidence that the PRLT feedback-P3 may be a state marker of alcohol dependence, informing us about when an individual stopped drinking alcohol following chronic use^36^. However, due to inconsistencies across samples and paradigms^17,25,34^ further research is needed to corroborate this finding.

One of the novel aspects of our work lies in using an unsupervised tensor decomposition algorithm to identify EEG patterns associated with alcohol dependence. Unlike traditional ERP methods, this data-driven approach makes no *a priori* assumptions, instead revealing the full range of features from population-level data^39^. We demonstrate using a combination of tensor components (R1-R2-R4) that we could predict group membership with a high classification accuracy of 80.4%. This model had exceptional performance when classifying HC, accurately predicting every participant using solely EEG data. By probing group differences in tensor component recruitment (Fig. 3B, Fig. 4C, Fig. 4D), we suggest that healthy non-dependent individuals, show an early focused parietal activity at 300 ms (R2) during reward valuation, which may be a neural signature of sensory information processing and evidence accumulation^40,41^. This process takes place before assessing the value-related features of the outcome, which are then used to update beliefs about reward contingencies, likely related to the subsequent centro-frontal activity peaking at 370 ms (R4)^11,12^.

Conversely, centro-frontal processing (R1) generally begins earlier in AD participants who exhibit this activation across the FRN and feedback-P3 windows (from 118 ms – 562 ms) and prior to parietal processing (R2). We hypothesise that R1, more actively recruited in short-term abstinence (Fig. 3E), represents early hyperfunctioning of centro-frontal circuits, potentially linked to an increase in sensitivity to value-related feedback^17,42^. Our model was less accurate at predicting which group AD participants belonged to, misclassifying 45% of AD as HC participants. This subset of AD had similar tensor component recruitment to the HC participants (Fig. 4C & 4D), and in general had a longer duration of abstinence. Notably, the temporally overlapping tensor components we uncovered here, would have been challenging to disentangle with ERP analysis: what appears as globally elevated signal in AD with ERPs (Fig. 2C, Fig. 2D) is resolved here into distinct temporal and spatial components (Fig. 3B).

Limitations of the study include challenges with participant recruitment due to lockdown restrictions imposed by COVID-19 and strict medication exclusion criteria which resulted in a smaller sample size than originally planned. Despite the small sample size, we demonstrate robust correlations following correction for multiple comparisons and tensor decomposition allowed us to uncover individual differences between participants. The sample of abstinent AD participants who were free from all psychiatric medication, may not be representative of the broader population, raising concerns about the generalisability of the findings. These exclusion criteria were however required to eliminate potential confounding factors, as commonly used medications in AD (e.g., selective serotonin reuptake inhibitors, naltrexone, and acamprosate) can also affect the brain’s reward system.

This study advances our understanding of the neural and behavioural adaptations in abstinence from alcohol dependence, showing EEG signal adaptation over time and improvements in learning deficits. Importantly, we demonstrate the power of unsupervised machine-learning to extract clinically meaningful EEG components from large amounts of unlabelled reward learning EEG data. Our findings could pave the way for low-cost EEG platforms in translational addiction psychiatry, facilitating advancements in screening of patients and at-risk populations^21^ and supporting novel treatment evaluation^22,23^. These platforms would focus on evaluation of the maladaptive decision-making circuity, which has emerged as a key target for novel interventions for addiction^14,15^.

## METHODS

### Participants

EEG data were recorded from 49 participants (including 28 HC and 21 AD) however 1 HC and 1 AD participant were excluded due to problems with EEG recording during the task. Furthermore, 1 HC participant was excluded due to clinically high anxiety (GAD-7 score > 15) which is known to impair probabilistic reversal learning and related EEG signal^43^. The analyses therefore include data from 26 HC participants (17 male) (mean age: 41, range: 24 - 60) and 20 abstinent AD participants (14 male) (mean age: 44.7, range: 31 – 58; mean abstinence: 20 months, range: 1 – 76 months) (Table 1). AD participants had been abstinent for at least 4 weeks, had severe alcohol dependency in the year preceding their current abstinence (i.e., DSM-V mean score: 10.75, score range: 9 - 11); and had never met the diagnostic criteria for any other substance use disorder^3^.

All participants had normal hearing, normal/corrected-to-normal vision and were fluent English speakers. HC participants had never met the DSM-V diagnostic criteria for alcohol or substance use disorder (excluding nicotine)^3^ and to their knowledge did not have any first-degree relatives with a history of alcohol or substance use disorder.

General exclusion criteria for all participants were as follows: (1) current primary axis I diagnosis (note: lifetime history of anxiety or major depressive disorders were permitted); (2) history of severe mental illness (e.g. bipolar, schizophrenia, affective disorder); (3) neurological illness (including but not limited to multiple sclerosis, stroke, epilepsy, space occupying lesions, Parkinson’s disease, vascular dementia, transient ischemic attack, clinically significant head injury); (4) Wechsler Adult Intelligence (WAIS) Intelligence quotient (IQ) score of below 70^44^, predicted using Wechsler Test of Adult Reading (WTAR) IQ score^45^ and demographic information (including age, gender, education history); (5) current use of psychoactive medication that would interfere with study integrity and cannot be stopped for the study duration (including but not limited to antipsychotics, anticonvulsants, antidepressants, disulfiram, acamprosate, naltrexone, varenicline); and (6) breast feeding and pregnancy.

Additional exclusions for each experimental session included positive urine drug test (note: AllTEST^TM^ 10 panel was used which includes tests for the cannabis, cocaine, opiates, amphetamines, benzodiazepines, tramadol, ketamine, methadone, methamphetamine and ecstasy) or positive alcohol breathalyser result. All participants were requested to refrain from recreational drug use in the 4 weeks prior to both experimental sessions.

Participants were recruited from Huddersfield and surrounding areas of Kirklees (UK) via posters, flyers, word of mouth, email campaigns and community Facebook groups. AD participants were also recruited from addiction recovery services (The Basement Project and CHART) and via a targeted Facebook campaign. This study was approved by the University of Huddersfield Department of Applied Sciences Ethics Committee (SAS-SREIC 1.5.19-1).

### Experimental sessions

All participants underwent a telephone screening to check basic eligibility criteria before attending a face-to-face clinical screening session (as detailed in^20^) which evaluated alcohol dependence/drug history, collected demographic, personality, and clinical information (Table 1) (Supplementary Methods). At the beginning of the clinical screening session participants gave written informed consent and completed a urine drug screen (AllTEST^TM^ 10 panel) and alcohol breathalyser. The researcher then conducted the DSM-V interview for drug and alcohol dependence, Mini-International Neuropsychiatric Interview ^46^, drug and alcohol timeline follow-back interview, Edinburgh Handedness Inventory (EHI) and Wechsler Test of Adult Reading (WTAR). Finally, participants were trained in the PRLT and completed 75 practice trials of the task.

Participants found to be eligible based on the clinical screening attended an EEG recording session. At the beginning of the session, they completed a urine drug screen (AllTESTTM 10 panel) and alcohol breathalyser. This was followed by 30 practice trials of the PRLT and the full EEG PRLT (including 300 trials, 21 minutes). Participants received £10 for attending the clinical screening, £15 for completing the EEG PRLT and were reimbursed for all travel costs. For the HC group, Amazon or supermarket vouchers were provided as reimbursement. For the AD group, only Amazon vouchers were provided to reduce risks associated with the visibility and immediate availability of alcoholic beverages in supermarkets.

### Probabilistic reversal learning task

The EEG PRLT included 3 blocks of 100 trials, each lasting approximately 7 minutes. At the end of each block, the participant had the opportunity to take a break and pressed the space bar when they were ready to begin the task again. To ensure participants were motivated, they were told that study payment would be based upon their task performance but were not informed how this would be calculated. In reality, all participants received the same fixed amount of £15 for their participation in the study. Before the start of each block, participants were shown three geometric cue symbols on the screen (e.g., A, B, C), these symbols were used in that block only and selected at random from a total set of 12 cue symbols. Participants were instructed to use trial and error to learn which cue symbol gave them reward feedback (i.e., a tick rather than a cross) most frequently (Fig. 1C). Throughout the task, one of the three cue symbols always had a 70% probability of giving tick feedback, i.e., the “high” reward probability symbol. The other two cue symbols had a 30% probability of giving tick feedback i.e., the “low” reward probability symbols. These exact probabilities were not disclosed to participants. The PRLT used here was adapted from^12^ and details of the task setup can be found in the Supplementary Methods.

## Modelling behavioural data

### Reinforcement learning (RL) models

#### Model 1

A classical Rescorla–Wagner (RW) model was used to compute trial-by-trial estimates of PE, *δ*(*t*), and expected value, *V*_*A*_(*t*), by considering the participants’ choices (i.e., selecting symbol A, B or C) and the feedback shown on each trial. The algorithm updated expected value on each trial using:

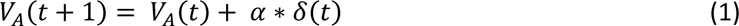

where *α* is the learning rate that controls how much the model adjusts the symbol’s expected value based on PE. The PE is calculated by the following equation: *δ*(*t*) = *r*(*t*) − *V*_*A*_(*t*), where *r*(*t*) denotes the feedback received on each trial (0 or 1, for negative or positive outcomes).

#### Model 2

An adapted Rescorla–Wagner learning model was implemented with an asymmetric learning model, with sperate learning rates for positive and negative outcomes, referred to here as the reward-punishment model (RP) model. This model considers that learning from oppositely valanced feedback may have different consequences on decision-making and behaviour^8,47^.

#### Model 3

A dynamic learning rate (DLR) model was implemented to model changes in task volatility (i.e., how stable and predictable the task was over time) for individual participants. This model includes a dynamic learning rate (*α*) which changes on a trial-by-trial basis based on updating of the choice’s expected value, as described in^12^ and summarised in the Supplementary Methods.

### Model fitting and model comparison

To find the best fitting model, we compared the different approaches (RW, RP, DLR) using a classical Bayesian Information Criterion (BIC) approach. We used the following equation to calculate BIC for each subject:

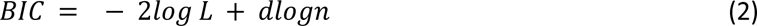

Where a complexity term (*dlogn*) scaled by the number of free parameters *d* and data points *n* (i.e., trials) is used to penalise goodness of model fit (− *log L*). BIC values were summed across all subjects and compared across models^12^.

### EEG data acquisition and pre-processing

EEG data acquisition and pre-processing followed the same procedures as detailed in^20^. EEG data were recorded at a sampling rate of 1000 Hz, using a 64-channel actiCAP with Ag/AgCl active electrodes in a 10-20 layout (Brain Products, Germany). Data were pre-processed with EEGlab Toolbox^48^ and Matlab R2019b. Raw continuous EEG data were downsampled to 500 Hz, high pass filtered at 0.1 Hz and sections with instructions/breaks were removed. Bad electrodes were rejected (Clean Rawdata EEGlab function) and interpolated, and 50 Hz noise was removed (CleanLine EEGlab plug-in version 1.04, https://www.nitrc.org/projects/cleanline). Independent component analysis (ICA) was used and artefact rejection was information by Multiple Artefact Rejection Algorithm (http://github.com/irenne/MARA). Data were low pass filtered at 40 Hz and re-referenced to TP9 and TP10. Feedback-locked epochs were extracted with a 500 ms prestimulus and 1500 ms poststimulus period, and a prestimulus baseline correction between −200 and 0 ms was applied to all epochs at each channel individually (ERPlab EEGlab plugin, https://erpinfo.org/erplab)^49^. Epoch rejection was not conducted thus preserving the entire dataset and maintaining statistical power^20,50^.

### Event Related Potential analyses

Based on seminal research defining the temporal dynamics of reward-based decision making in healthy subjects^13^, the FRN and feedback-P3 were measured as the mean amplitude over 180 – 260 ms and 280 - 380 ms following feedback onset, respectively. All ERPs were quantified over a central cluster of electrodes (FC1, FCz, FC2, C1, Cz, C2, CP1, CPz and CP2)^13^.

Group differences were evaluated using mixed model analyses of covariance (ANCOVA) with age and IQ included as covariates. To evaluate valence processing, 2 x 2 ANCOVA models used trial outcome (Prediction Error is defined as PE: i.e., tick symbol = positive-PE and cross symbol = negative-PE) as the within-subjects factor and group (i.e., HC group, AD group) as the between-subjects factor. To evaluate magnitude processing, 2 x 2 ANCOVA models used trial PE (i.e., high-PE and low-PE) as the within-subjects factor and group (i.e., HC group, AD group) as the between-subjects factor. Based on trial-by-trial PE calculated using the best fitting RP model, the top 15% and bottom 15% of trials in terms of PE were extracted for high-PE and low-PE conditions, respectively (giving a total of 45 trials per condition). These tests were performed separately for the FRN and feedback-P3.

Mauchly’s Test of Sphericity was used to check the assumption of sphericity in ERP data submitted to the ANCOVA models. Greenhouse-Geisser correction was applied where violation occurred. R software (version 4.0.5) was used for these analyses and a significance level of *p* < 0.05 was applied.

### Tensor Decomposition analyses

The feedback-locked epochs (i.e., trials) (K) were reduced to the main timepoints (T) of interest for feedback processing (-100 and 700 ms) and then collated across all participants (P) and electrodes (E) into a 4-way data tensor *X*^(4)^ ∈ ℝ ^∈ × *E* × *P* × *K*^, where P = 46, T = 401, E = 62 and K = 299. A tensor decomposition model based on NCP with a hierarchical alternating least squares method was applied to the 4-way data tensor using the MATLAB Nonnegative Matrix and Tensor Factorization Algorithms Toolbox^51^ and Tensor Toolbox^52^. Since the group averaged ERPs for the two main components of interest in this task (FRN and feedback-P3) have amplitudes (µV) with positive values (Fig. 2A) a non-negative constraint was applied to values within the factor matrices (**A, B, C, D, E**) to give the low approximation **<u>X</u>** ∼ **A** ◦ **B** ◦ **C** ◦ **D** ◦ **E** with 4 outer product tensor components (i.e., Rank (R) = 4) (Fig. 1B)^53^. Furthermore, non-negativity has the benefit of yielding sparser and largely non-overlapping representations of the input data, which enhances interpretability of the identified components^28,54,55^.

Based on the outer product (◦) decomposition, scalar tensor entries *X*_*tepk*_ can be estimated with:

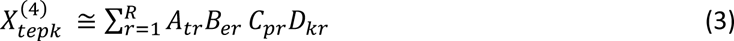

Where R, the tensor rank, is the number of outer product components deemed to adequately explain the data. Matrix *A* ∈ ℝ ^*T* × *R*^ is the temporal factor and its columns 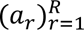 represent the level of activation for each component at each time point. Matrix *B* ∈ ℝ ^*E* × *R*^ is the spatial factor and its columns 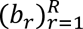 represent the electrode weightings for each component. Matrix *C* ∈ ℝ ^*P* × *R*^ is the participant factor and *D* ∈ ℝ ^*K* × *R*^ is the trial factor which captured the level of activation of each component for each participant and on each trial, respectively.

To select the number of components R, we evaluated the trade-off between complexity of the output, data approximation and robustness of the extracted components. Varying the number of components, we empirically found that R=4 yielded physiologically plausible EEG components that captured hypothesised group differences in EEG responses and that increasing the number of components did not alter these main spatiotemporal patterns or substantially reduce the relative error of the tensor decomposition model.

Two-sample t-tests were used to evaluate differences in the level of recruitment for each tensor component between the between the HC and AD group. One test was completed for each of the four tensor components with values extracted from the columns 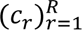 of the participant factor matrix *C* ∈ ℝ ^*P* × *R*^ which represented the level of recruitment of each component for each participant. Tests which were significant (*p* < 0.05) following FDR correction (4 tests) are included in the main results.

### Classifying groups via Support Vector Machine

We aimed to predict participant groups from the tensor components using a Medium Gaussian support vector machine (SVM) classifier trained using leave-one-out cross validation, chosen based on suitability for the small sample size and implemented with MATLAB’s Classifier Learner app. Values extracted from the columns 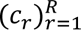 of the participant factor matrix *C* ∈ ℝ ^*P* × *R*^ were used as predicators of the HC and AD group. An iterative process was used to identify participant factors and combinations thereof which were most predictive of participant group. Firstly, the individual factors were entered into the classifier separately (R1, R2, R3, R4), then pairs (R1-R2, R1-R3, R1-R4, R2-R3, R2-R4, R3-R4), then triplets (R1-R2-R3, R1-R2-R4, R1-R3-R4, R2-R3-R4) and finally all factors together^53^. Classifier performance was evaluated based on percentage of accurately classified participants i.e., classifier accuracy and area under the ROC curve (AUC).

### Investigating the effects of abstinence on EEG signal and behaviour

Firstly, mixed effects linear models were conducted within the AD group to evaluate associations between ERP data and abstinence duration. We performed eight regression models, with the ERP mean amplitude (µV) as the dependent variable: FRN positive-PE, FRN negative-PE, FRN high-PE, FRN low-PE, P3 positive-PE, P3 negative-PE, P3 high-PE, P3 low-PE. Independent variables included abstinence duration, total units of alcohol consumed per adult year of drinking, IQ and age. Only models in which an independent variable was significantly associated with a dependent variable following FDR correction (8 tests) are included in the results (*p* < 0.05). The equation for these regression models is as follows:

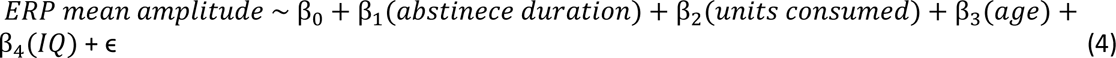

Secondly, correlations were used to explore whether abstinence duration in the AD group was associated with: (1) main performance measure from the PRLT (i.e., learning reversals); (2) the IQ of participants, or (3) recruitment of the first tensor component (R1), since this component was more commonly recruited among AD compared to HC participants. The Shapiro–Wilk test confirmed abstinence duration was not normally distributed; therefore, Spearman correlation was used for all tests and the FDR corrected p-values are reported (i.e., 3 tests) implemented using the p.adjust function in R.

Finally, to understand whether classification of participants using tensor decomposition components was influenced by abstinence, a two-sample t-test was used to compare abstinence duration between misclassified and correctly classified AD participants within the highest performing classification model. R software (version 4.0.5) was used for all statistical analyses.

## ACKNOWLEDGEMENTS

This research was supported by a UKRI Medical Research Council Future Leaders Fellowship grant and BBSRC grant (MR/T023007/1 and BB/Y001494/1) (to E.F.F.), BBSRC grant (BB/Y513799/1) (to I.D) and the NIHR Manchester Biomedical Research Centre (NIHR203308) (to M.K). The views expressed are those of the author(s) and not necessarily those of the NIHR or the Department of Health and Social Care. Funding provided by the University of Huddersfield funded the PhD studentship that supported data collection (M.K). We wish to thank Change Grow Live and the Basement project for promoting the project. The views expressed are those of the author(s) and not those of CGL and associated services. Finally, we wish to thank Oliver Young, Ihtisham Ahmed, Katie West, Aisha Lunat and Micheala Marsden for assistance with EEG data collection.

## CONFLICT OF INTEREST

The authors declare that they have no known competing financial interests or personal relationships that could have appeared to influence the work reported in this paper.

## AUTHORSHIP CONTRIBUTION

M.K: Conceptualization, Data curation, Investigation, Formal analysis, Project administration, Writing – original draft. C.R: Methodology, Writing – review & editing. A.M: Writing – review & editing. I.D: Conceptualization, Methodology, Investigation, Formal analysis, Supervision, Writing – review & editing, Visualization. E.F: Conceptualization, Methodology, Formal analysis, Supervision, Visualization, Writing – original draft, review & editing.

## Notes

### Competing Interest Statement

The authors have declared no competing interest.

### Summary of Updates

I would like to change the title so it can be linked with the version which may be published Altered EEG markers of reward learning during abstinence in alcohol dependence: a probabilistic reversal learning study

