## Supplementary Information for "Altered EEG markers of reward learning during abstinence in alcohol dependence: a probabilistic reversal learning study"

### Experimental Sessions

The questionnaire collected demographic information plus a comprehensive battery of clinical questionnaires which were hosted by the online Qualtrics platform including:

- Alcohol Use Disorder Identification Test (AUDIT)^1^, a short 10-item self-rating tool to screen for alcohol harm and unhealthy alcohol use (e.g., risky or hazardous consumption or any alcohol use disorder), in which AD participants were instructed to complete questions 1-3 relating alcohol consumption based on their drinking behaviours in the year prior to their current period of abstinence;
- Obsessive Compulsive Drinking Scale (OCDS), a short 14-item self-rating tool that measures an individual's alcohol use and their attempts to control drinking, in which Questions 7 - 10 relating to current drinking behaviours were excluded as all AD participants were currently abstinent;
- Generalised Anxiety Disorder Assessment (GAD-7), a 7-item tool that is used to assess the severity of generalised anxiety disorder based on symptoms present over the last 2 weeks^2^;
- Beck Depression Inventory (BDI), a 21-item self-report tool used to assess severity of depression^3^; and
- Childhood Trauma Questionnaire (CTQ), a 28-item self-rating tool which measures severity of emotional abuse and neglect, physical abuse and neglect and sexual abuse^4^, which all participants were given opportunity to opt out of answering this questionnaire to avoid triggering post-traumatic stress relating to childhood trauma.

### Probabilistic reversal learning task

**Task setup & stimuli**

The task included twelve abstract geometric cue symbols (examples are shown in Fig. 1A) and two feedback symbols (a tick for positive feedback and a cross for negative feedback). All stimuli were presented in black on a grey background and equivalent luminance and contrast including the cue (150 × 150 pixels), feedback (200 x 200 pixels) and fixation (30 × 30 pixels) symbols. Presentation software (Neurobehavioral Systems Inc., Albany, CA) was used for task programming and stimuli were presented at 60 Hz framerate, on a Windows 10 computer (64 bit, 3GB RAM, nVidia GeForce GT 710 graphics card) with a monitor resolution of 1920 x 1080 pixels. Participant were sat 50 cm away from the monitor and responded by pressing the left or right arrow buttons on a USB keyboard using the index and middle finger on their dominant hand.

**PRTL detailed paradigm**

At the start of each trial a fixation cross was shown in the centre of the screen for a random jitter of 1– 1.2 seconds. Next, two out of the three cue symbols were then shown on either side of the central fixation cross for a maximum of 1.25 seconds. Across the trials the three cue symbols were displayed in a fixed order using three different pair combinations (i.e., trial 1 = AB, trial 2 = BC, trial 3 = CA) and their position on the screen (left or right) was randomised. Participants were informed about this design to prevent them from attempting to learn non-existent patterns. Participants indicated which symbol they believed to be the “high” reward probability symbol, by selecting either the left or right keyboard arrow for the left and right symbols, respectively. Once the keyboard arrow had been selected, the fixation cross flickered for 0.1 seconds to indicate their selection had been recorded. After a random jitter of 1– 1.2 seconds, feedback for that trial was shown on the screen for 0.75 seconds. Their choice of symbol was either rewarded or unrewarded by giving the participant a tick or a cross, respectively. If the participant failed to press the left or right arrow within the 1.25 seconds a “Lost trial” message was shown on the screen and no other feedback was given.

Once the participant had learnt through trial and error which cue symbol was the one with “high” reward probability, a reversal occurred, meaning “high” reward probability was switched to a different cue symbol chosen at random from the other two used in that block. Participants were deemed to have learnt the rule when they consistently selected the “high” reward probability symbol in five out of the previous six trials, this learning criterion ensured reversals were only experienced after adequate learning. Buffer trials were included once the learning criterion was met to reduce the predictability of reversals. A Poisson process was used to determine the number of buffer trials (from 1 up to a maximum of 8). The design maintained a balanced number of trials with tick and cross feedback and forced participants to make a choice between the two low probability symbols even after they had learnt the rule. This was important because it meant participants were shown an equal number of rewarded versus unrewarded trials^5^.

### Modelling behavioural data

**Dynamic learning rate**

The learning rate for each trial $\left( i \right)$ is adjusted by the slope of the smoothed absolute PE ($m$) (a measure of how much the difference between the expected reward and the actual reward changes over time) using these update equations:

$\alpha\left( i \right)=\alpha\left( i-1 \right)+ f\left( m \left( i \right) \right).\left( 1- \alpha\left( i-1 \right) \right), if m> 0_{2}$	(1)

$\alpha\left( i \right)=\alpha\left( i-1 \right)+ f\left( m \left( i \right) \right).\left( \alpha\left( i-1 \right) \right), if m< 0$	(2)

In this context, $f\left( m \left( i \right) \right)$represents a double sigmoid function that converts the slope (m) into a range between 0 and 1. This function adjusts how much the learning rate changes from one trial to the next. Importantly, it's controlled by a free parameter (*γ*). When *γ* is high, the adjustments to the learning rate become so small that it practically remains constant, acting as if the learning rate is fixed^5^.

**Fixed parameters across models**

For all models evaluated:

(1) a SoftMax decision function determined the probability of selecting a specific symbol in each trial ($t)$, based on the values of the symbols on the screen:

$P_{A}\left( i \right)= \sigma\left( \beta\left( V_{A}\left( t \right)- V_{B}\left( t \right) \right)-\varphi\right)$	(3)

Here, $\sigma(z) = 1/(1 + e^{-z})$is the logistic function, $\varphi$ is the point where choosing each option is equally likely, and β measures how random or consistent the choices are (i.e., the exploration/exploitation parameter).

(2) The initial values for all symbols (A, B and C) were set to 0.33 and initial learning rates set to 0.4.

(3) Only the value of the symbol selected by the participant to have “high” reward probability was updated on each trial, the expected value of the unselected symbol and the symbol not shown on the screen for that trial were not updated.

### Classifying alcohol and control participants from EEG data

**Supplemental Table 1.** Performance of Medium Gaussian Support Vector Machine models tested using combinations of tensor components (Acc = Accuracy, AUC = Area Under Area Under ROC Curve, FPR = False Positive Rate, TPR = True Positive Rate)

| Tensor Component Included in Model | Acc | AUC | FPR vs TPR |
| --- | --- | --- | --- |
| R1 | 76.1% | 0.66 | 0.08, 0.55 |
| R2 | 60.9% | 0.55 | 0.15, 0.30 |
| R3 | 56.5% | 0.19 | 0.00,0.00 |
| R4 | 56.5% | 0.60 | 0.54, 0.70 |
| R1 & R2 | 76.1% | 0.75 | 0.08, 0.55 |
| R1 & R3 | 73.9% | 0.70 | 0.12, 0.55 |
| R1 & R4 | 76.1% | 0.71 | 0.08, 0.55 |
| R2 & R3 | 54.3% | 0.51 | 0.27, 0.30 |
| R2 & R4 | 63.0% | 0.69 | 0.27, 0.50 |
| R3 & R4 | 54.3% | 0.55 | 0.50, 0.60 |
| R1 & R2 & R3 | 71.7% | 0.75 | 0.12, 0.50 |
| R1 & R2 & R4 | 80.4% | 0.73 | 0.00, 0.55 |
| R1 & R3 & R4 | 71.7% | 0.71 | 0.15, 0.55 |
| R2 & R3 & R4 | 65.2% | 0.61 | 0.23, 0.50 |
| R1 & R2 & R3 & R4 | 76.1% | 0.71 | 0.04, 0.50 |
